# Chemokines form complex signals during inflammation and disease that can be decoded by extracellular matrix proteoglycans

**DOI:** 10.1101/2022.09.20.508420

**Authors:** Amanda JL Ridley, Yaqing Ou, Richard Karlsson, Nabina Pun, Holly L Birchenough, Thomas A Jowitt, Craig Lawless, Rebecca L Miller, Douglas P Dyer

## Abstract

Chemokine driven leukocyte recruitment is a key component of the immune response and is central to a wide range of diseases. However, there has yet to be a clinically successful therapeutic approach that targets the chemokine system during inflammatory disease; possibly due to the supposed redundancy of the chemokine system. A range of recent studies have demonstrated that the chemokine system is in fact based on specificity of function. Here we have generated a resource to analyse chemokine gene (ligand and receptor) expression across different species, tissues and diseases; revealing complex expression patterns whereby multiple chemokine ligands that mediate recruitment of the same leukocyte type are expressed in the same context, e.g. the CXCR3 ligands CXCL9, 10 and 11. We use biophysical approaches to show that CXCL9, 10 and 11 have very different interactions with extracellular matrix glycosaminoglycans (GAGs) which is exacerbated by specific GAG sulphation. Finally, *in vivo* approaches demonstrate that GAG-binding is critical for CXCL9 driven recruitment of specific T cell subsets (e.g. CD4^+^) but not others (e.g. CD8^+^), independent of CXCR3 expression. Our data demonstrate that chemokine expression is complex and that multiple ligands are likely needed for robust leukocyte recruitment across tissues and diseases. We also demonstrate that ECM GAGs facilitate decoding of these complex chemokine signals so that they are either primarily presented on GAG-coated cell surfaces or remain more soluble. Our findings represent a new mechanistic understanding of chemokine mediated immune cell recruitment and identify novel avenues to target specific chemokines during inflammatory disease.

## Introduction

Leukocyte migration and recruitment facilitates tissue inflammation which in turn is central to a plethora of diseases such as rheumatoid arthritis, inflammatory bowel disease and cancer(*1*). Leukocyte recruitment is itself primarily driven by chemokines (chemotactic cytokines), making them key players in inflammatory based disease and prime therapeutic targets(*2*). Chemokines are a large family of small proteins that are thought to function by binding to their concomitant receptors on circulating leukocytes(*3*). This interaction produces integrin activation, enabling firm adhesion of leukocytes to the blood vessel wall and subsequent trans-endothelial migration.

Despite their importance the chemokine system has yet to be therapeutically targeted. The reasons for this are multiple, however, a central problem has been the idea of redundancy where multiple chemokine ligands bind to multiple receptors and multiple receptors bind to multiple ligands(*2, 4*). Several recent studies have set out to determine whether redundancy is a key aspect of chemokine function and have largely demonstrated the opposite, i.e., extreme specificity of the chemokine system(*5–10*). This can be understood from a receptor perspective as recent studies have demonstrated that receptor expression is fine-tuned to have specific receptors on the leukocyte cell surface at each stage of its migration to facilitate specific functions(*9*).

In contrast, when an unbiased analysis is undertaken during inflammatory scenarios a range of different chemokines with over-lapping functions are present in the same location(*8*). This presents a challenge to the idea of specificity of how non-redundant migratory outcomes are produced when multiple ligands for the same receptor are present in the same environment. One mechanistic explanation for this is biased agonism, where different ligands can produce different functional outcomes via the same receptor via different interactions(*11, 12*). However, this does not fully explain how these specialised functions of chemokines are produced since the differing receptor affinities should in theory dominate which ligands are bound to a receptor at any given time. One way that differential localisation and receptor-binding availability may be achieved is via interactions with extracellular matrix (ECM) glycosaminoglycans (GAGs)(*10, 13*).

ECM GAGs are particularly present within the glycocalyx that lines the luminal endothelial surface within blood vessels but can also be found throughout tissues(*14*). Chemokine:GAG interactions have been shown to be key for leukocyte recruitment *in vivo* as they facilitate endothelial retention of chemokines on the endothelium in the presence of blood flow(*15, 16*) and in the case of CXCL4 may directly mediate its function(*17*). However, specific comprehension of the role of chemokine:GAG interactions on the cell surface and within tissues in leukocyte recruitment is lacking.

Here we show that chemokine ligands and receptors are present in complex, but distinct, families across tissues and disease. These families contain chemokine ligands with over-lapping abilities to recruit the same leukocytes. In particular, the CXCR3 ligands CXCL9, 10 and 11 display over-lapping expression patterns across a wide range of tissues and diseases and have differential roles in recruitment of CXCR3^+^ cells. We also demonstrate that these ligands have very different interactions with ECM GAGs. These differential interactions facilitate regulation of ligand localisation at the cell surface or in solution. We also found that these interactions play surprisingly specific roles in CXCL9 mediated leukocyte recruitment.

## Results

### Chemokine ligands and receptors form complex and over-lapping signals

To better understand how chemokines co-ordinate the immune response via leukocyte recruitment we undertook a systematic view of the transcriptional relationship between chemokine ligands and their receptors during inflammation and disease. To do so we analysed transcriptomic data that has been deposited in the EMBL-EBI expression atlas (Fig. 1)(*18*). The database was downloaded followed by minimal “cleaning” of the data to deliberately take an unbiased approach to analysis (Supp. Fig. 1). Output data was then probed using principal component analysis (PCA) and correlation matrix analysis to reveal transcriptional relationships between chemokine ligands and receptors across a range of tissues and diseases in humans and mice (Table 1).

**Figure 1.**
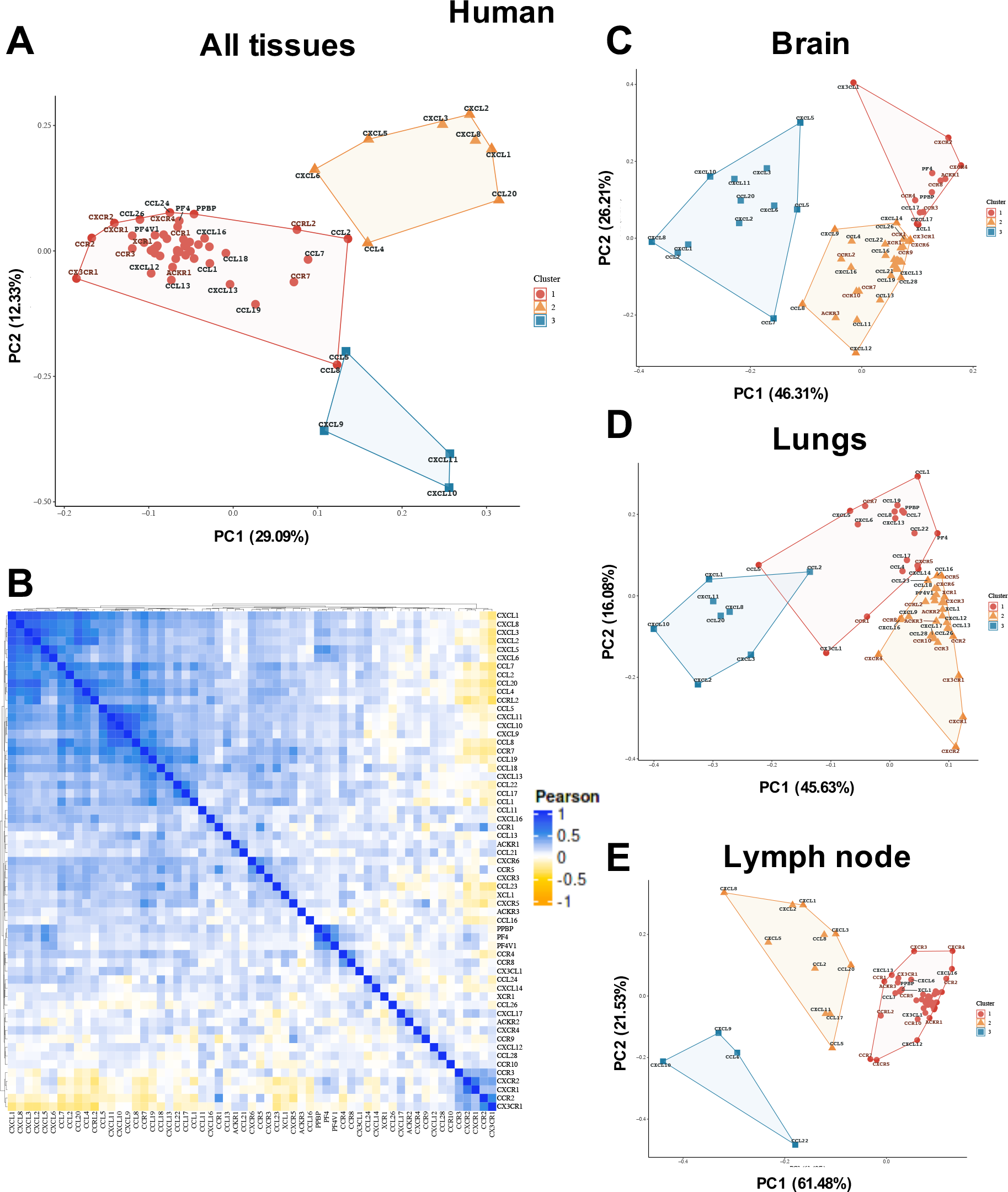
Chemokine receptors and ligands are present in complex and tissue specific patterns during inflammation. The EMBL-ELI expression atlas was analysed for relatedness on expression of all chemokine ligands and receptors in human data from (A) PCA analysis, (B) heat map analysis of all data pooled or separated into PCA analysis of (C) brain, (D) lungs or (E) lymph node.

**Table 1.** Flow cytometry antibodies to surface and intracellular markers used to characterize murine samples.

| Antigen | Fluorochrome | Clone | Manufacturer | Working Dilution |
| --- | --- | --- | --- | --- |
| CXCR3 | BV421 | CXCR3-173 | BioLegend | 1:200 |
| IgG Isotype | BV421 | HTK88 | BioLegend | 1:200 |
| CD4 | FITC | GK1.5 | BD Biosciences | 1:200 |
| CD8 | PerCP/Cy5.5 | 53-6.7 | BioLegend | 1:800 |
| F4/80 | APC | BM8 | eBioscience | 1:200 |
| Ly6C | Af700 | HK1.4 | BioLegend | 1:200 |
| Ter119 | APC/ef780 | TER-119 | eBioscience | 1:200 |
| CD3 | APC/ef780 | 17A2 | eBioscience | 1:100 |
| TCRβ | APC/ef780 | H57-597 | eBioscience | 1:200 |
| Ly6G | BV510 | 1A8 | Biolegend | 1:100 |
| CD11c | BV605 | N418 | BioLegend | 1:200 |
| B220 | BV650 | RA3-6B2 | BioLegend | 1:100 |
| CD11b | BV711 | M1/70 | BioLegend | 1:800 |
| CD45 | BV785 | 30-F11 | BioLegend | 1:200 |
| CD64 | PE | X54-5/7.1 | BioLegend | 1:100 |
| SiglecF | PE/CF594 | E50-2440 | BD Biosciences | 1:400 |
| NK1.1 | PE/Cy5 | PK136 | BioLegend | 1:200 |
| TCR$\gamma\delta$ | PE/Cy7 | eBioGL3 (GL-3,<br>GL3) | eBioscience | 1:200 |

Immediate patterns emerge from this analysis in both the human (Fig. 1 and Supp. Fig. 2) and mouse (Supp. Fig. 3) data, firstly separating into 3 distinct clusters. The largest group, in both the human (cluster 1, Fig. 1A and B) and mouse (cluster 2, Supp. Fig. 3) data, contains a range of genes that have a diverse function in the recruitment of different leukocytes. The other two groups, however, contain genes with closely related functions. The first contains chemokine ligands and receptors that are primarily associated with recruitment of neutrophils are present, specifically CXCL1, CXCL2, CXCL3, CXCL5, CXCL6 and CXCL8 (human cluster 2) and CXCL1, CXCL2, CXCL3, CXCL5 and CCL2 (mouse cluster 1)(*3*). These clusters also contain some additional genes usually associated with recruitment of other cell types, e.g., CCL4 (monocyte) and CCL20 (T cell) in the human data set and CCL2, CCL3, CCL4 and CCL7 (monocyte) in the mouse dataset. Pointedly in both the human and mouse dataset cluster 3 contains CXCL9, CXCL10 and CXCL11 which all signal through CXCR3, primarily in the recruitment of T cells(*19*).

These data demonstrate that chemokine gene expression is associated with specific phases of the immune response so that those genes recruiting the same types of leukocyte are often similarly expressed.

### Individual tissues and diseases have specific chemokine ligand and receptor expression patterns

Next we analysed patterns of chemokine ligand and receptor expression more closely by separating the data into different tissues, focussing on the brain (Fig. 1C), lung (Fig. 1D) or lymph node (Fig. 1E). The human (Fig. 1C) and mouse (Supp. Fig. 3) brain data demonstrated clustered expression of neutrophilic chemokines. Human brain cluster 3 contains CXCL1, CXCL2, CXCL3, CXCL5 and CXCL8. Mouse cluster 1 was limited to CXCL1 and CXCL2 alongside a number of monocytic chemokines, suggesting species-specific inflammatory responses in the brain. The clusters in the human lung (Fig. 1D), mouse lung (Supp. Fig. 3C), human lymph node (Fig. 1E), and mouse lymph node (Supp. Fig. 3D) were tissue-specific, but again the close relatedness of the neutrophilic chemokines was maintained.

We next separated the data into diseases for which there were at least ten entries in the expression atlas, namely Crohn’s disease, ulcerative colitis, psoriasis, rheumatoid arthritis and Alzheimer’s disease (Fig. 2). As above, specific signatures were seen in each context with the first four diseases containing a neutrophilic chemokine cluster. In contrast, the gene relatedness signatures appeared very distinct in the Alzheimer’s disease analysis compared to the others, suggesting the chemokine system is functioning differently in this context.

**Figure 2.**
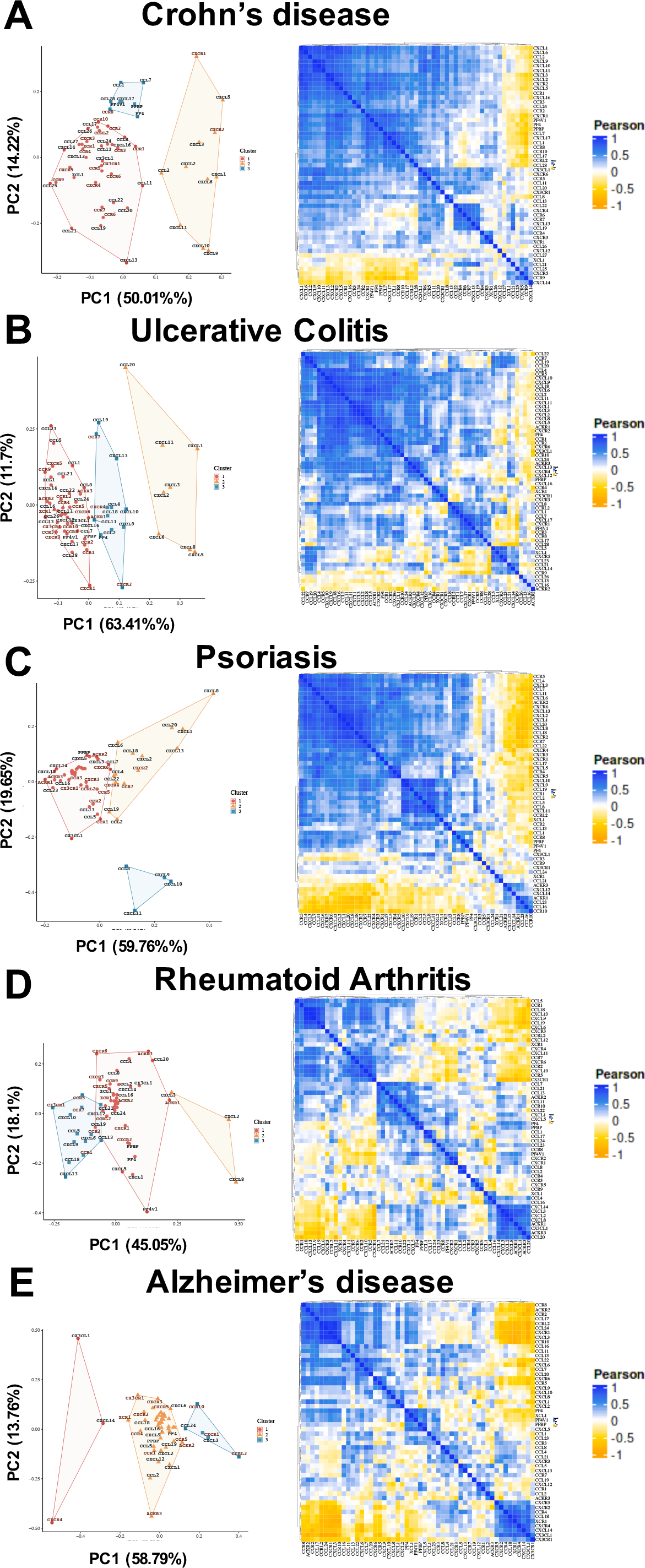
Chemokine receptors and ligands are present in disease specific patterns. The EMBL-ELI expression atlas was analysed for relatedness on expression of all chemokine ligands and receptors in human data from (A) Crohn’s disease (B) ulcerative colitis, (C) psoriasis, (D) rheumatoid arthritis and (E) Alzheimer’s disease.

The separation of genes into related clusters was observed across the separated tissues and diseases with specific signatures, reflecting the differential nature of the inflammatory response between tissues and disease. However, certain patterns were clear across these situations, with strikingly consistent clustering of a range of chemokines that can all recruit neutrophils. Strikingly the clustering of CXCL9, CXCL10 and CXCL11 appeared to be present across the separated tissues and diseases.

### CXCR3 ligands are consistently expressed together across tissues and disease

Our data thus present a potential example of supposed chemokine redundancy, where CXCL9, 10 and 11, which all bind and signal through CXCR3 (Fig. 3A), may be consistently co-expressed across different inflamed tissues and diseases. To determine the degree of this co-expression we specifically analysed the expression relatedness between the receptor CXCR3 and its ligands CXCL9, 10 and 11 (Fig. 3B and C). This approach confirmed that CXCL9, 10 and 11 are very closely related in expression during inflammation across a range of tissues and diseases. In contrast, though closely related to each other these ligands have a much lesser transcriptional relationship with their receptor CXCR3 (Fig. 3), suggesting that the ligands and their receptor are usually produced at distinct sites.

**Figure 3.**
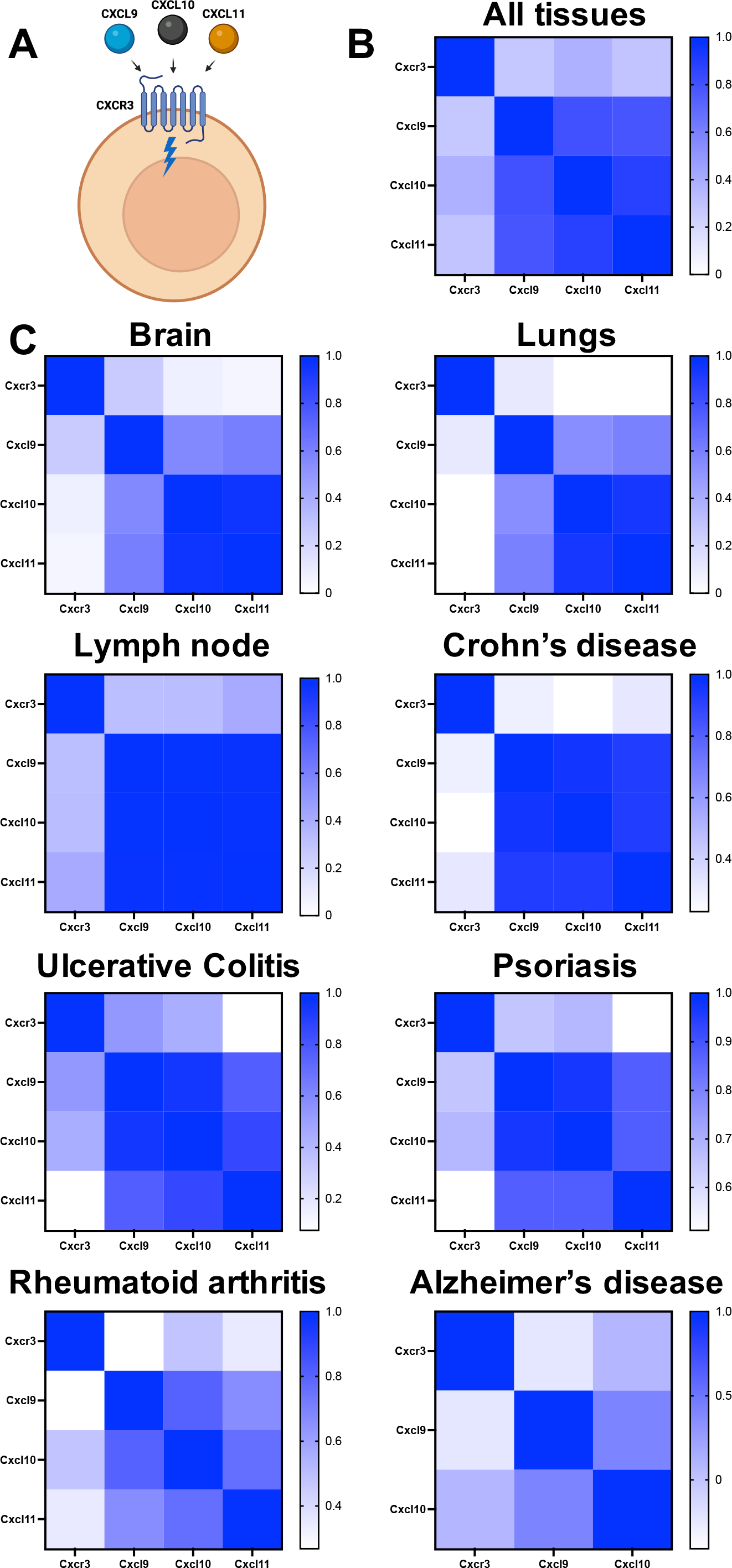
The CXCR3 ligands CXCL9, 10 and 11 are expressed in over-lapping patterns across tissues and disease. (A) CXCL9, 10 and 11 can all bind and signal through the chemokine receptor CXCR3 that is typically found on T cells. (B) The EMBL-ELI expression atlas (human) was analysed for relatedness in expression of CXCR3 and its’ ligands CXCL9, 10 and 11 across all tissues or (C) in distinct tissues and diseases.

This close transcriptional relatedness between ligands but not their receptor is also evident in other families, for example, CXCR1 and 2 and their ligands, where CXCR1 and 2 are also closely transcriptionally related, demonstrating their co-operative function in co-ordinating leukocyte recruitment (Supp. Fig. 4). In contrast, CXCR5, CXCR6, CCR1, CCR4, CCR6, CCR10 are more closely transcriptionally related to their ligands (Supp. Fig. 4-6). Whilst CCR7 and CCR8 have a close transcriptional relationship to one ligand (CCL19 and CCL1, respectively) but not the other (Supp. Fig. 6). These differing relationships may reflect function in local leukocyte positioning, where receptors and ligands would be closely transcriptionally related, versus long range leukocyte recruitment, where they would not be closely related.

### CXCL9, 10 and 11 have specific functions *in vivo*

The numerous examples of overlapping expression of CXCL9, 10 and 11 suggests that each is required and thus plays a specific role during the recruitment of CXCR3^+^ cells. We next confirmed this relatedness at the protein level *in vivo* in the mouse carrageenan inflamed air pouch recruitment model (Fig. 4A and B). The air pouch *in vivo* leukocyte recruitment model was then chosen to determine whether the CXCR3 ligands have a differential function in leukocyte recruitment *in vivo*. We injected equimolar amounts of CXCL9, 10 and 11 into the air pouch (Fig. 4A), allowing analysis of a wide range of leukocytes (Fig. 4C and D and Supp. Fig. 7 and 8). None of the ligands produced statistically significant changes in overall (CD45^+^) leukocyte recruitment (Fig. 4E) or in the number of neutrophils, macrophages, or eosinophils (Supp. Fig. 9). However, CXCL9 did produce a significant increase in the number of T cells (TCRβ^+^) in contrast to CXCL10 and 11 (Fig. 3E) (flow cytometric gating strategy Supp. Fig. 7 and 8). This demonstrates that CXCL9, 10 and 11 do not have the same effect on leukocyte recruitment *in vivo*, supporting previous findings that they each play a specific role in recruitment of T cells(*11, 19*).

**Figure 4.**
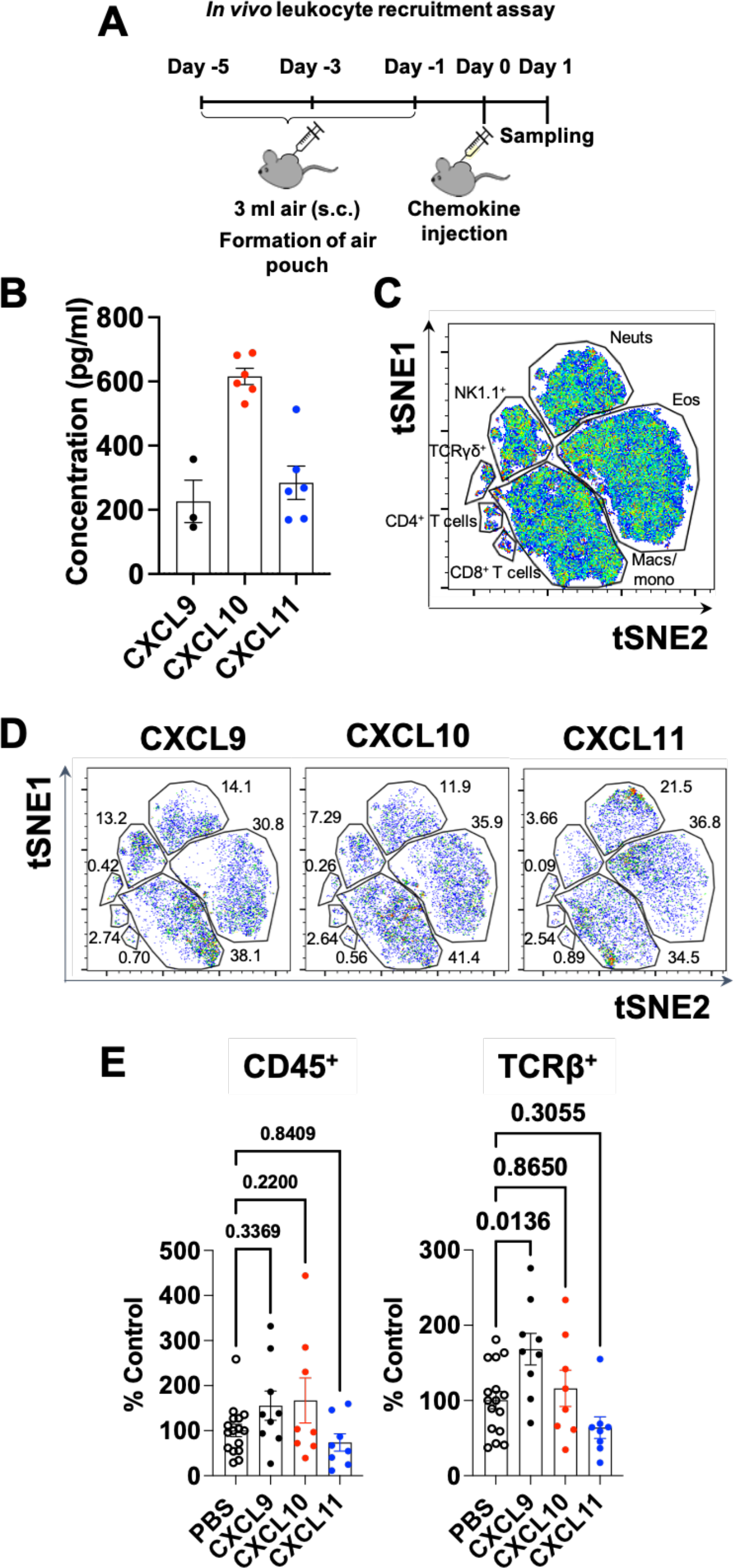
CXCL9, 10 and 11 have different abilities to mediate cell recruitment *in vivo*. (A) Schematic of the *in vivo* air pouch leukocyte recruitment model. (B) Analysis of chemokine concentration in the carrageenan inflamed air pouch. (C) Representative tSNE of all murine cells gated on live, single, CD45^+^ and built on CD4, CD8, F4/80, Ly6C, Ter119, CD3, TCRβ, CXCR3, Ly6G, CD11c, B220, CD11b, CD64, Siglec F, NK1.1 and TCRγδ. FlowSOM clusters are illustrated by gates. (D) tSNE analysis of air pouches injected with equimolar amounts of CXCL9, 10 and 11. (E) Quantification of all leukocytes (CD45^+^) and T cells within the air pouch following injection of CXCL9, 10 or 11. E analysed using a one-way ANOVA.

### Interactions with ECM GAGs decodes CXCL9, 10 and 11 signals

The data above demonstrate that CXCL9, 10 and 11 are produced in over-lapping combinations and play different roles in T cell recruitment *in vivo.* Therefore, we next sought to understand how these complex chemokine signals could be decoded to allow each to play its specific biological role. We hypothesised that differential interactions with GAG side chains on ECM proteoglycans (Fig. 5A) may produce differential localisation of these ligands within the ECM and on the cell surface.

**Figure 5.**
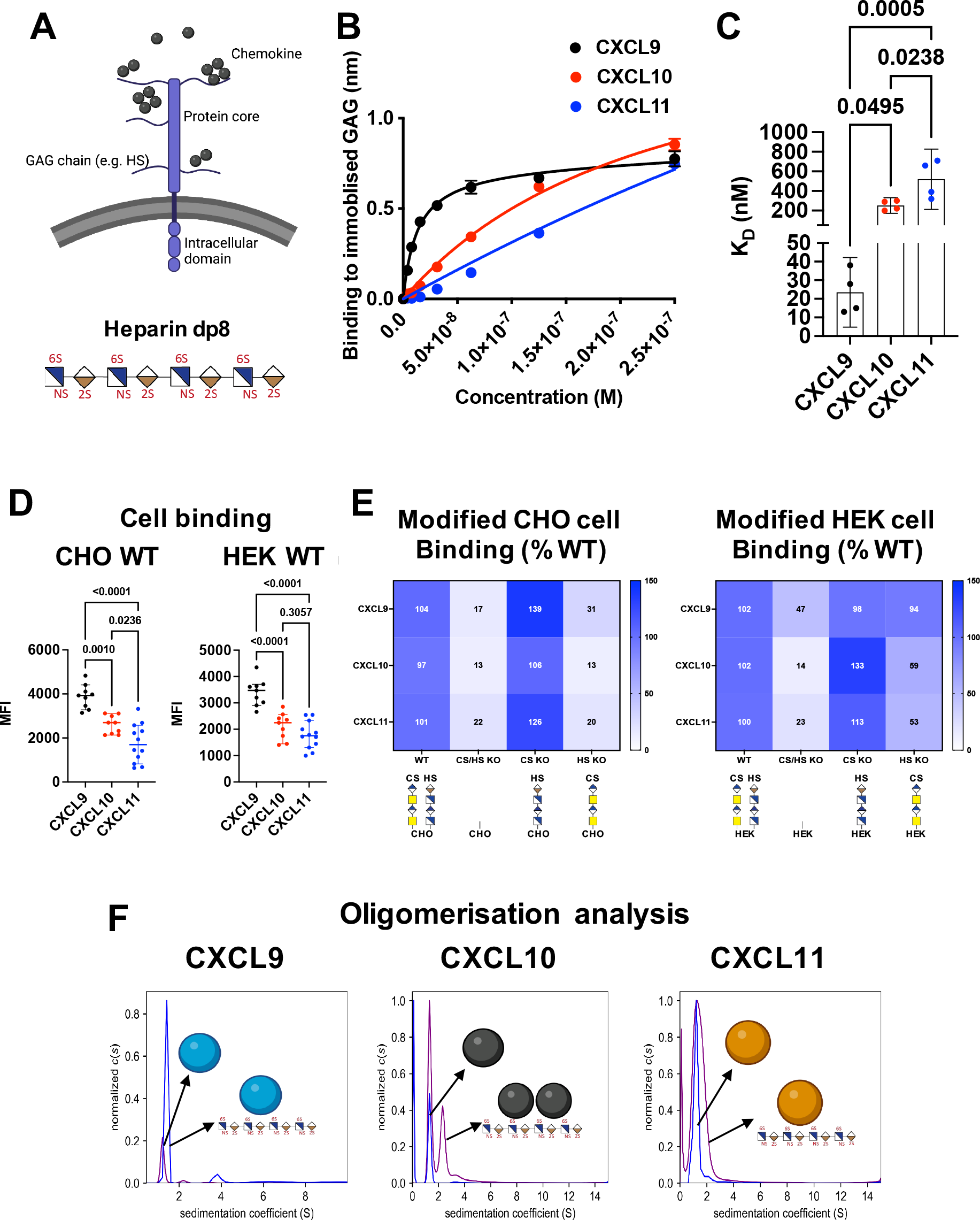
CXCL9, 10 and 11 have different affinity interactions with ECM GAGs. (A) GAGs are sugar side chains found on cell surface and ECM proteoglycans and purified dp8 can be used as a model GAG. (B) Immobilised dp8 binding to chemokines in BLI equilibrium signal is plotted against chemokine concentration. (C) BLI data in B was analysed for steady state affinity (*K*_D_) estimates. (D) Binding of biotin-labelled chemokines to WT CHO and HEK293 cells quantified by mean fluorescent intensity (MFI) detected using flow cytometry. (E) Binding of biotin-labelled chemokines to genetically modified CHO cells with KO of B4galt7 (CS/HS KO), CSgalnact1/2/Chsy1 (CS KO) and Extl3 (HS KO), and HEK293 cells with KO of B4GALT7 (CS/HS KO), CHSY1/3 (CS KO), and EXTL3 (HS KO). All data was normalized to MFI for WT. (F) AUC analysis of chemokine oligomerization state in the absence and presence of heparin dp8 GAG. Data plotted ± SEM, (B) is representative of two separate experiments, (C) is pooled data from two separate experiments, (D) contains data from three separate pooled experiments, (E) is the mean of three separate pooled experiments and (F) is representative of two separate experiments. C and D analysed using a one-way ANOVA.

We utilised bio-layer interferometry (BLI) biophysical analysis to study the interaction between these ligands and isolated GAG sugar models (heparin dp8). BLI demonstrates that CXCL9 (24 ± 12 nM), CXCL10 (253 ± 50 nM) and CXCL11 (520 ± 194 nM) have significantly different GAG affinity estimates (Fig. 5B and C). To analyse chemokine:GAG interactions in a cellular context, binding of labelled CXCL9, CXCL10, and CXCL11 to WT Chinese hamster ovary (CHO) or human embryonic kidney (HEK) cells was performed using flow cytometry (Fig. 5D). In both cases the order of binding (demonstrated by MFI signal) was CXCL9>CXCL10>CXCL11, in agreement with the BLI studies.

To determine which GAGs are responsible for this cellular chemokine binding we used genetically engineered cells that express distinct types of chondroitin sulfate (CS) and heparan sulfate (HS) GAG chains(*20, 21*). We first compared chemokine binding to WT, CS/HS knock out (KO), CS KO, and HS KO cells derived from CHO and HEK293 cell lines (Fig. 5E). CXCL9, 10 and 11 all bound to HS on CHO cells since its removal reduced binding, but neither bound to CS since binding was not reduced in its absence. CXCL10 and CXCL11 bound to HS on HEK cells, in contrast CXCL9 binding is only reduced when both HS and CS are removed from HEK cells, suggesting that CXCL9 can bind to both in a potentially redundant fashion. Oligomerisation has been shown to be key to differential chemokine:GAG interactions (*22, 23*). Therefore, we next determined the differential oligomerisation of CXCL9, 10 and 11 in the absence and presence of the heparin dp8 GAG using analytical ultracentrifugation (AUC) (Fig. 5F). All three primarily exist as monomers when alone in solution. In the presence of dp8 (ratio 1:2, chemokine:GAG) CXCL9 remains primarily monomeric, CXCL10 becomes 50% dimeric and CXCL11 remains primarily monomeric.

Together these results show that CXCL9, CXCL10 and CXCL11 have a very different ability to bind and be retained on cell surface proteoglycans and will thus be differently distributed within a tissue or on cell surfaces *in vivo*.

### GAG fine structure facilitates specificity of binding to CXCL9, CXCL10 and CXCL11

GAG sulphation has previously been shown to drive interaction specificity with other chemokines, e.g., CCL2 and CXCL4(*17, 24*). We, therefore, hypothesised that GAG sulphation points could produce further differentiation in binding to CXCL9, 10 or 11. GAG sulphation patterns are primarily produced during GAG synthesis by a wide range of different sulfotransferases. Acting on HS are the NDSTs, HS2ST1, HS6STs and HS3STs responsible for N−, 2-O, 6-O, and 3-O sulphation respectively (Fig. 6A)(*25*). Together these modifications can achieve incredible sequence specificity.

**Figure 6.**
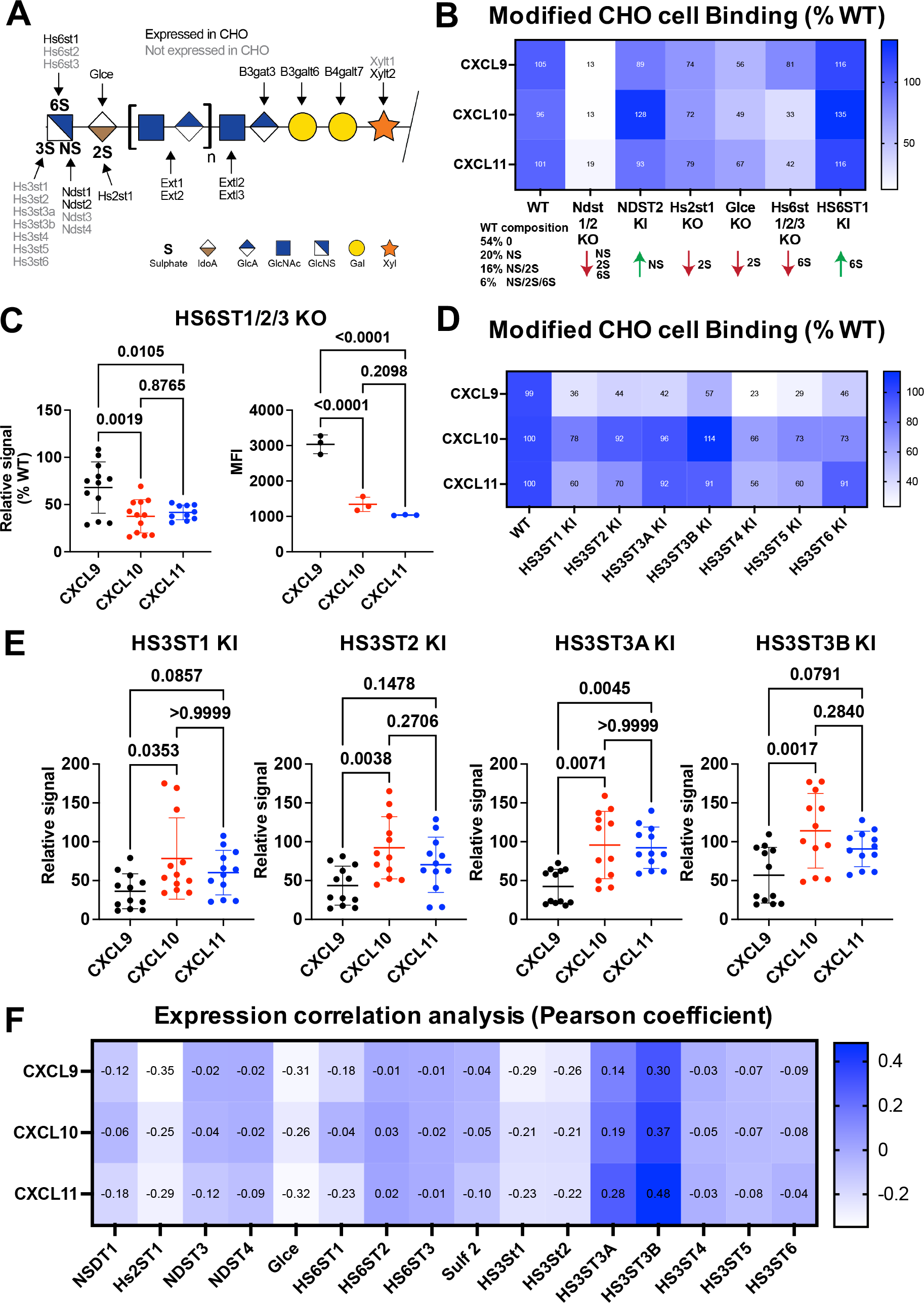
ECM GAG sulphation mediates differentiation of binding to CXCL9, 10 and 11. (A) Schematic of HS GAG structure, including sulphation points and the enzymes that produce them. (B) Normalised (relative to wild type) binding of labelled CXCL9, 10 or 11 to genetically modified CHO cells. (C) Normalised and <u>absolute</u> binding of CXCL9, 10 and 11 to CHO cells in which KS^ST1/2/3 have been genetically removed. (D and E) Normalised binding of CXCL9, 10 and 11 to CHO cells genetically engineered to express the enzymes regulating 3-O GAG sulphation. (F) EMBL-ELI expression atlas analysis of relatednessCXCL9, 10 and 11 and GAG sulphation gene expression. B and D, data plotted as mean from three separate pooled experiments. C and E data plotted as mean ± SEM from three separate pooled experiments and analysed using a one-way ANOVA.

In order to dissect their differential contribution to CXCL9, 10 and 11 binding we utilised the recently developed GAGOme cell library, which is a large panel of CHO cells genetically engineered to display distinct GAG features on the cell surfaces (Fig. 6A)(*20, 26*). Flow cytometry studies revealed that the biggest contributor to HS binding in this context for all three ligands was N-sulfation by NDST1 and 2, unsurprising given that this is the first step in the biosynthetic pathway and essential for priming and modification of different sulfation patterns (Fig. 6B). The effect of HS2ST1 (2-O sulphation) and GLCE (epimerase enhancing levels of 2-O sulphation) removal reduced binding comparably across all three ligands. Combined removal of HS6ST1/2/3 (6-O sulphation) had a much greater effect in reduction of relative binding to CXCL10 and CXCL11 than CXCL9 (Fig. 6B). When absolute rather than relative signal is analysed this difference is exacerbated, in the absence of HS 6-O sulphation the cell surface GAGs have 3 times the capacity to bind CXCL9 compared to CXCL10 and CXCL11 (Fig. 6C).

Using the GAGOme approach we also determined the relative effect of 3-O sulphation of HS on binding by using CHO cells in which the different HS3ST isoenzymes responsible for this modification have been knocked-in (KI) (Fig. 6D and E). Adding in 3-O sulphation largely reduced CXCL9 binding, had no significant effect in CXCL10 binding and in some cases reduced CXCL11 binding. There is again evidence of specificity; HS3ST1, HS3ST2, HS3ST3A and HS3ST3B KI cells all have significantly lower binding signal for CXCL9 compared to CXCL10 and CXCL11.

Given the ability of the GAG sulphation genes to regulate differential binding to chemokines we next sought to determine whether there was a transcriptional relationship between them. We analysed the human expression atlas database from all human tissues pooled for the different GAG synthesis and sulphation genes (Supp. Fig. 10-12). Specifically, analysis of the transcriptional relationship between CXCL9, CXCL10, CXCL11 and the different enzymes that facilitate HS sulphation suggests no positive correlation with the genes facilitating N−, 2-O or 6-O sulphation (Fig. 6F). This may suggest that the chemokine ligands are produced at distinct times/locations from these sulphation enzymes. In contrast, analysis of the transcriptional relationship between the 3-O sulphation genes and CXCL9, 10 and 11 reveals reasonable correlation of transcription between HS3ST3A and HS3ST3B and these chemokine ligands (Fig. 6F). This suggests that these genes can be expressed at the same time/location and may collaborate to produce specific presentation of CXCL9, CXCL10 and CXCL11.

Global comparison for the GAG synthesis, modification and proteoglycan protein core genes demonstrated less discrete clustering than is the case with chemokine ligand and receptor genes when analysing pooled tissues and diseases (Supp. Fig. 11 and 12). However, more discrete clusters become apparent when the data is separated into specific tissues in both humans and mice.

These data show that in addition to the general interactions with GAGs specific sulphation types can add an additional layer of specificity to these interactions to facilitate differential geographical localisation of ligands that bind to the same receptor.

### Interactions with ECM GAGs have a specific role in CXCL9 function *in vivo*

Given the ability of ECM GAGs to bind to CXCL9 and preferentially retain it on the cell surface we next sought to determine whether this interaction is important to its’ function *in vivo* (Fig. 6). We first determined which of the cells would be recruited by CXCL9 due to their CXCR3 expression (Fig. 7A and Supp. Fig. 7 and 8). As expected, CXCR3 was present on a number of T cell subsets. Pre-incubation of CXCL9 with purified GAG, to block interaction with endogenous cell surface GAG, inhibited CXCL9 mediated recruitment of CD4^+^ T cells and possibly TCRγδ cells, but not NK cells or CD8^+^ cells (Fig. 7B).

**Figure 7.**
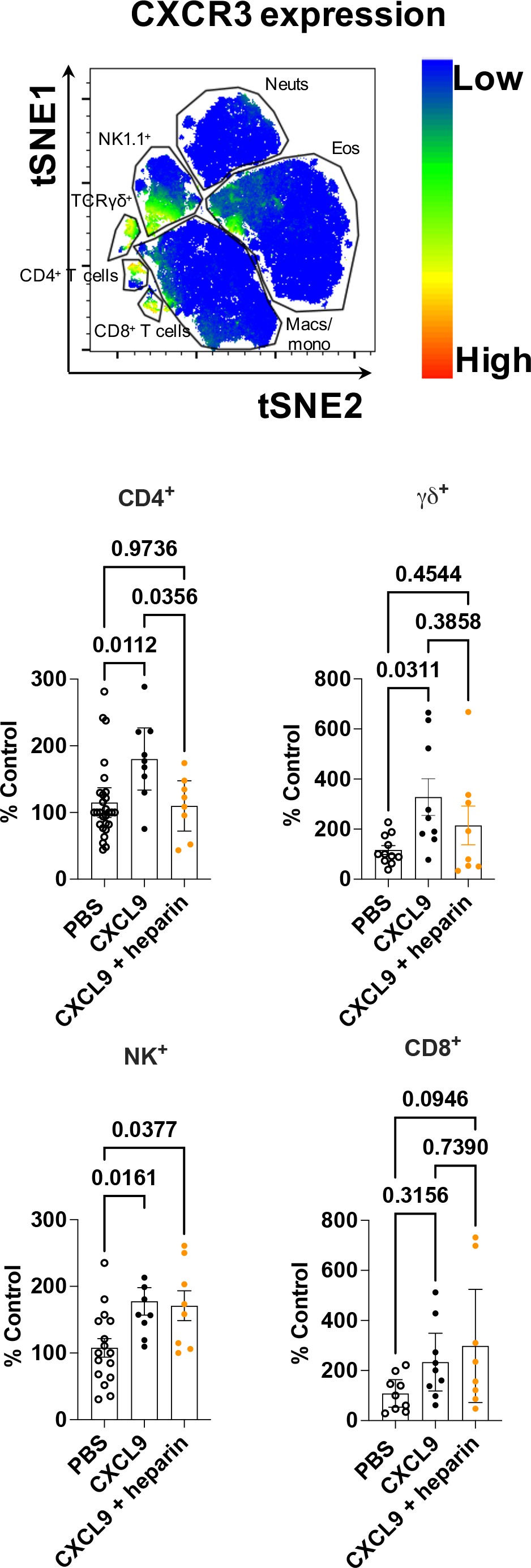
CXCL9:GAG interaction plays specific roles in mediating recruitment of different CXCR3^+^ cells. (A) Representative tSNE analysis of CXCR3 expression of chemokine recruited leukocytes gated on live, single, CD45^+^ and built on CD4, CD8, F4/80, Ly6C, Ter119, CD3, TCRβ, CXCR3, Ly6G, CD11c, B220, CD11b, CD64, Siglec F, NK1.1 and TCRγδ. (B) Normalised leukocyte counts of air pouches injected with the indicated solutions. (B) Data expressed ± SEM from three pooled separate experiments analysed using a one-way ANOVA.

These surprisingly specific effects of blocking CXCL9 binding to endogenous GAG suggests that the interaction between CXCL9 and cell surface GAGs plays a vital yet differential role in the recruitment of individual CXCR3^+^ T cells *in vivo*.

## Discussion

Here we show that complex signals of chemokines with over-lapping function can be decoded by ECM GAGs to facilitate specific chemokine localisation during inflammatory disease (Fig. 8). We hypothesise that this differential localisation is critical to the specific function of individual chemokine ligands that has been demonstrated in recent studies(*10*). Alongside these studies, our data may further challenge the classic theory of chemokine redundancy that has been thought to preclude therapeutic targeting of the chemokine system in inflammatory disease(*10*).

**Figure 8.**
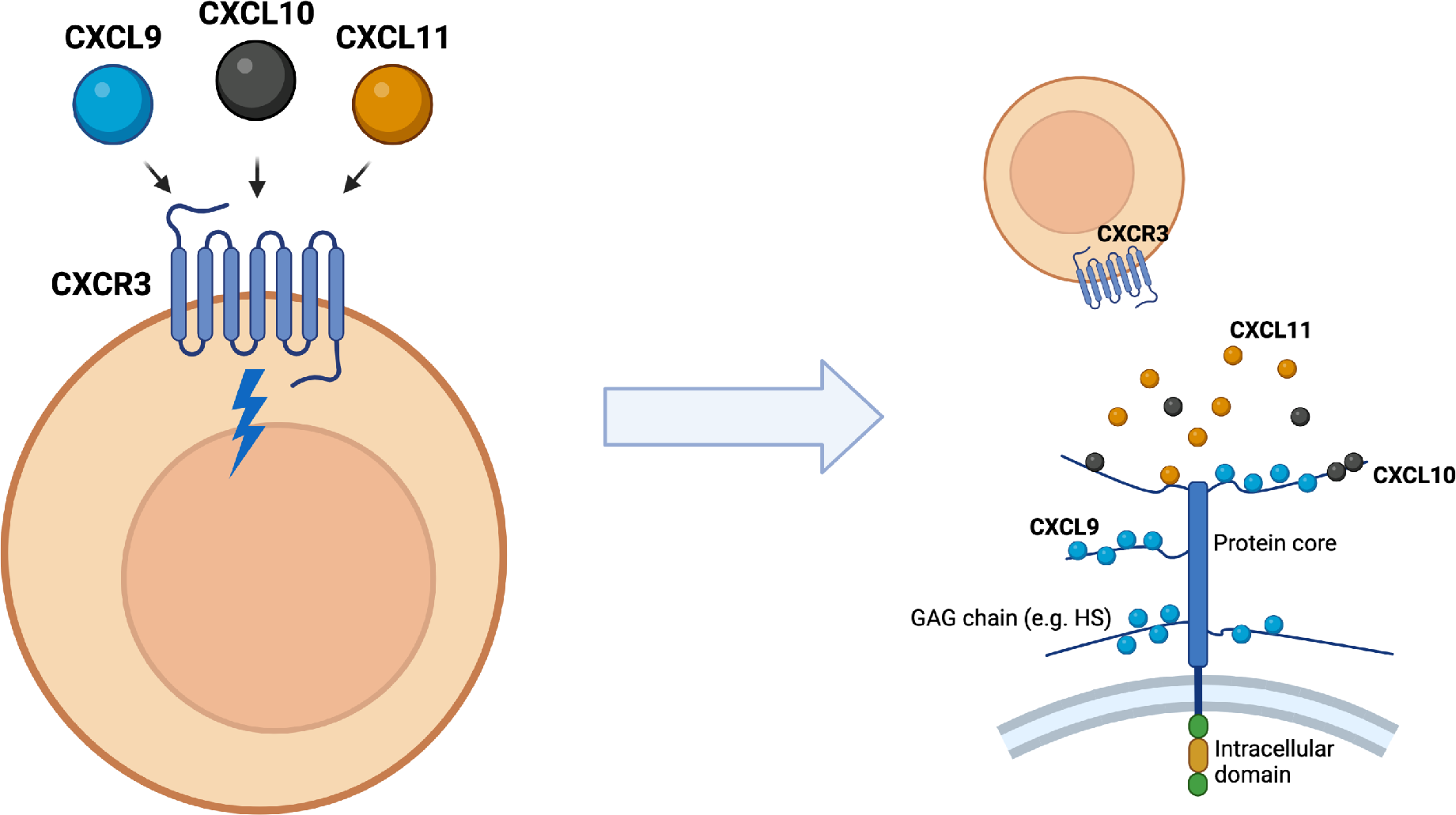
Complex chemokine signals are “decoded” via interactions with ECM GAGs. (A) CXCL9, 10 and 11 all bind to the same receptor with different affinities and biased signalling outcomes and are found in over-lapping expression patterns during inflammation and disease. (B) Differential GAG interactions means that CXCL9 is more likely to be retained on GAGs on the cell surface or within the ECM, with CXCL10 and CXCL11 being more likely to be present in their soluble state.

Whilst individual papers have presented data showing the presence of multiple chemokine ligands that recruit the same leukocyte, we are not aware of previous unbiased and comprehensive analyses to better define the expression relationship between all chemokine ligands and receptors. Our findings of specific clusters of chemokines and receptors with over-lapping function confirm assumptions that are made about the immune response, which previously have little data to support them. Specifically, given that the early inflammatory response is primed to recruit neutrophils it may not be surprising that we consistently find distinct clusters across tissues and diseases that contain chemokines associated with neutrophil recruitment. Similarly, the consistent overlapping production of the CXCR3 ligands, CXCL9, 10 and 11 may also be unsurprising as there will be distinct stages of the immune response where CXCR3^+^ T cells are required to fight infection/disease. The fact that in multiple instances chemokines are produced at the same time with supposedly redundant functions supports the idea that each actually performs a specific function in leukocyte recruitment and that multiple chemokine ligands are required to mediate recruitment of a given cell type.

The data we present herein of consistently overlapping expression of the CXCR3 ligands presents a fundamental problem to our understanding of chemokine biology. Namely, how would a migrating CXCR3^+^ T cell be able to interpret a complex signal containing all 3 ligands in the same environment to achieve recruitment and positioning? Our data evidencing the role of ECM GAGs in mediating differential immobilisation of these chemokines on ECM GAGs on the cell surface, and potentially within tissues, may solve this problem. Our data may also explain any potential differences in GAG-mediated protection of chemokines from proteolytic degradation(*27*) and support the idea of targeting GAGs to inhibit CXCL9 function in disease (*16, 28, 29*). Overall our data suggest that a CXCR3^+^ cell is unlikely to encounter these ligands in the same geographical location even with their close transcriptional relationship due to differential GAG binding. This also overcomes the problem whereby if all 3 ligands were present in their soluble form then CXCL11 would dominate binding to CXCR3 due to its much higher affinity for the receptor compared to CXCL9 and 10(*11*).

Furthermore, our data revealed that GAG sulphation can add additional layers of specificity to the chemokine:GAG interaction with 6-O sulphation particularly differentiating HS interactions with CXCL9, 10 and 11. The role of HS 3-O sulphation in biology is much less understood than for N−, 2-O or 6-O sulphation due to the problems associated with its analysis(*30*). Strikingly we found that 3-O sulphation mediates differentiation in binding to the CXCR3 ligands and its presence actually reduces binding to CXCL9. Excitingly these data generate the hypothesis that cells and/or tissues may tune their sulphation pattern, e.g., during inflammation, on their ECM GAGs to selectively bind and present certain chemokines over others.

Our study enhances the wider understanding of the co-ordinated role of the chemokine system during inflammation and disease across specific tissues. We also reveal a role for ECM GAGs in decoding the complex chemokine signals that are produced in these contexts to facilitate the specificity of the chemokine system during leukocyte recruitment.

## Materials and methods

### Materials

Up to 4 mice were housed in cages of up to four in a 12 hr light/dark cycle, with free access to food and water. All experiments were carried out following ethical approval from The University of Manchester and University of Glasgow and under licence from the UK Home Office (Scientific Procedures Act 1986). All chemokines were purchased from Protein Foundry and dp8 and heparin GAGs were purchased from Iduron.

### Bioinformatic Analysis

#### Dataset Construction

To explore gene co-expression, complete microarray and RNA-sequencing differential expression results were downloaded from Expression Atlas in December 2020(*18*). In total, 807 projects (2,450 assays) that operated in *Homo sapiens* and 894 projects (2,431 assays) that operated in *Mus musculus* were selected. Then, accessory description files were used to filter out all tumour-related projects and identify related tissues. In this study, brain, lung, spleen and lymph node were selected.

#### Assay Integration

According to literature and preliminary studies, we selected 136 genes that potentially play important roles in chemokine activities during the immune response (*3*). Those 136 genes were categorised into five groups: chemokine ligands (44 genes), chemokine receptors (23 genes), matrix (glycosaminoglycan) synthesis genes (17 genes), matrix metalloproteinase (24 genes), and proteoglycan synthesis (28 genes).

All analytics results of these 136 genes from tumour-free assays were then collected and concatenated together into one data frame. If the research was forced on a specific organ, only relevant assays were selected. Later, a four-step filtration was carried out to minimise the missing value of each assay but to keep more genes in the data. Firstly, genes that were missed in more than 70% of assays were removed. Secondly, in the output data frame from the last step, assays that contained less than 90% of genes were filtered out. Then, all assays with missing values were cleared. The final output data frame would then be ready for further investigations.

#### Co-expression Exploration

In order to figure out co-expression relationships between chemokine-related genes, correlation matrixes and principal component analysis (PCA) were plotted. In the correlation matrix, Pearson correlation coefficients between each pair of genes were calculated and visualised as a heatmap using Complex Heatmap package in R (Version 4.1.0)(*31, 32*). PCA plots were drawn using autoplot package(*33*). Gene cluster results are computed by Clustering Large Applications (CLARA) algorithm(*34*). All plots and correlation tables are available at https://shiny.its.manchester.ac.uk/mqbpryo2/ChemoInt.

### *In vivo* leukocyte recruitment assay analysis

#### Air pouch formation

Air pouches were formed by three subcutaneous injections of 3 ml of sterile air under the dorsal mouse skin every 48 hrs. One day after the final injection the chemokine or carrageenan (1% w/v, Simga-Aldrich) was injected into the air pouch. 24 hrs after injection mice were culled and the air pouch flushed with PBS containing 1% FCS and 1 mM EDTA (two occasions of 3 ml) and analysed for cellular and protein content as below.

#### Flow cytometry analysis

Post cell isolation, cells were plated at 1 × 10^6^ cells per well in v-bottom Nunclon Delta treated 96-well plates (Thermo Fisher Scientific) and washed twice with 100 μl ice-cold 1X PBS to remove proteins in the supernatant before being pelleted by centrifugation at 500 × *g* for 2 min at 4 °C. Cells were then incubated with a Live/Dead amine reactive viability dye (Zombie ultraviolet (UV) dye) (BioLegend) diluted 1:2000 in 10 μl 1X PBS for 15 min at RT in the dark to facilitate dead cell exclusion. Incubation steps were performed in the dark to prevent fluorochrome bleaching.

To prevent non-specific binding of antibodies via Fc receptors on the cell surface, all samples were incubated with FcR block (5 μg ml^−1^ αCD16/CD32 (2.4G2; BD Biosciences)) in 50 μl flow buffer (PBS containing 1% FCS) for 10 min at 4 °C. After blocking, cells were centrifuged at 500 × *g* for 2 min at 4 °C before pelleted cells were resuspended in 50 μl flow buffer containing surface marker antibodies at the dilutions shown in Table 1 for 30 min at 4 °C. Following incubation, cells were washed in 150 μl flow buffer and pelleted by centrifugation at 500 × *g* for 2 min at 4 °C. This wash step was then repeated with a further 200 μl flow buffer. After washing, cells were resuspended in 1% paraformaldehyde (PFA) (Sigma-Aldrich) for 10 min at RT in the dark to prevent the dissociation of antibodies from their target molecules. Cells were then centrifuged at 500 × *g* for 2 min at 4 °C before being resuspended in 200 *μ*l flow buffer ready for acquisition. Cells were stored at 4 °C prior to analysis. In some instances, after blocking, cells were incubated with CXCR3 (1:200) or an IgG isotype control (1:200) in 50 μl flow buffer for 15 min at 37 °C before being washed twice and stained with surface marker antibodies as described above.

To calculate cell counts in some experiments, CountBright Absolute Counting Beads (Thermo Fisher Scientific) were used. Before acquisition of BAL samples on the flow cytometer, 15 μl beads were added per sample. During analysis, beads were identified based on their high side scatter (SSC) and low forward scatter (FSC) phenotype. To calculate absolute cell count, the total number of beads added along with the number of bead events acquired was compared to the volume of cells added and total number of cell events acquired, as per the manufacturers protocol.

tSNE and FlowSOM analysis were performed in R (Version 4.1.3) using CD4, CD8, F4/80, Ly6C, Ter119, CD3, TCRβ, CXCR3, Ly6G, CD11c, B220, CD11b, CD64, Siglec F, NK1.1 and TCRγδ.

#### ELISA and Luminex analysis of air pouch fluid

Air-pouch fluid contents were obtained as described in (*8*). Specific concentrations of CXCL9 were measured by enzyme-linked immunosorbent assay (ELISA), using the mouse CXCL9 ELISA kit (R&D Systems) in a 96-well high binding ELISA plate following the manufacturer’s instructions. Plates were read on a VersaMax Microplate Reader (Marshall Scientific) at 450 nm. The samples were also analysed using a Bio-Plex Pro Mouse Chemokine Panel, 31- Plex Assay (Bio-Rad, UK). Samples were read and the data was acquired on a Bio-Plex Manager™ (Software version 6.2).

### Chemokine:GAG interaction analysis

#### Bio-layer interferometry (BLI)

An Octet Red96 system (Sartorius AG, Goettingen, Germany) was used with a methodology adapted from (*22*). GAGs had biotin attached at their reducing end with a previously described approach (*35*) before immobilisation to High Precision Streptavidin (SAX) biosensors (Sartorius AG, Goettingen, Germany). To achieve this SAX biosensors were hydrated for 10 mins in assay buffer (10 mM Hepes, 150 mM NaCl, 3 mM EDTA, 0.05% Tween-20, pH 7.4). Immobilisation of heparin dp8 GAG (0.078 *μ*g/ml) was done in assay buffer until an immobilisation level of approx. 0.1 nm was reached. Sensors were subsequently washed with regeneration buffer (0.1 M Glycine, 1 M NaCl, 0.1% Tween, pH 9.5) and re-equilibrated in assay buffer. Blank reference or GAG coated sensors were then dipped into 200 *μ*L of assay buffer containing chemokines at the indicated concentrations for at least 180 sec (association) before being transferred to assay buffer containing wells (dissociation) for at least 180s before a regeneration buffer wash step. The binding signal was recorded throughout and the signal from binding of chemokine to blank (no immobilised GAG) sensors and by GAG immobilised sensors in assay buffer alone was subtracted. As well as a signal over time, the maximum signal during the association phase of the interaction was recorded and used to generate *K*_D_ value esimates in the Octet analysis software. Data were acquired at 5 Hz and analysed using the Octet HT 10.0 analysis programme.

#### Analytical ultra-centrifugation

CXCL9, CXCL10 or CXCL11 were re-suspended in PBS to a final concentration of 50 *μ*g/ml either alone or in the presence of heparin dp8 at a ratio of 1:2 (chemokine:GAG). Samples were loaded into 2-sector cells with PBS as a reference and centrifuged at 50,000 rpm in a 4-hole An60Ti rotor monitoring the absorbance at 230 nm until sedimentation was reached. The time-resolved sedimenting boundaries were analysed using Sedfit (*36*). The resulting profiles are shown in Gussi (*37*).

#### Cell surface GAG binding

This analysis was performed as previously (*20*), using libraries of CHO Glutamine synthetase (GS) −/− or HEK293 6e cells genetically engineered by CRISPR/Cas9 for KO and Zinc finger nucleases for KI of genes(*20, 21, 26*). 1 × 10^5^ cells were washed in PBS before incubation with biotinylated recombinant human CXCL9, CXCL10 or CXCL11 (10 *μ*g/mL) (Protein Foundry LLC) in assay buffer (PBS + 1% FBS) for 30 min at 4 °C. Cells were washed with assay buffer followed by incubation with Alexa Fluor 488-streptavidin (1:2000 in assay buffer) (S32354, Invitrogen) for 30 min at 4 °C. Cell were again washed and then re-suspended in assay buffer and fluorescence intensity was analysed using a SA3800 spectral cell analyzer (SONY). All experiments were performed a minimum of three times using triplicate samples.

### Statistics

Statistical analysis were performed using Prism (GraphPad). Experiments containing two groups were analysed using an unpaired T-test and data containing more than two groups were analysed using a one-way ANOVA with a post-hoc multiple comparison test. *P* < 0.05 was considered to be statistically significant.

