## Supplementary figures for "Chemokines form complex signals during inflammation and disease that can be decoded by extracellular matrix proteoglycans"

Supplementary Figure 1

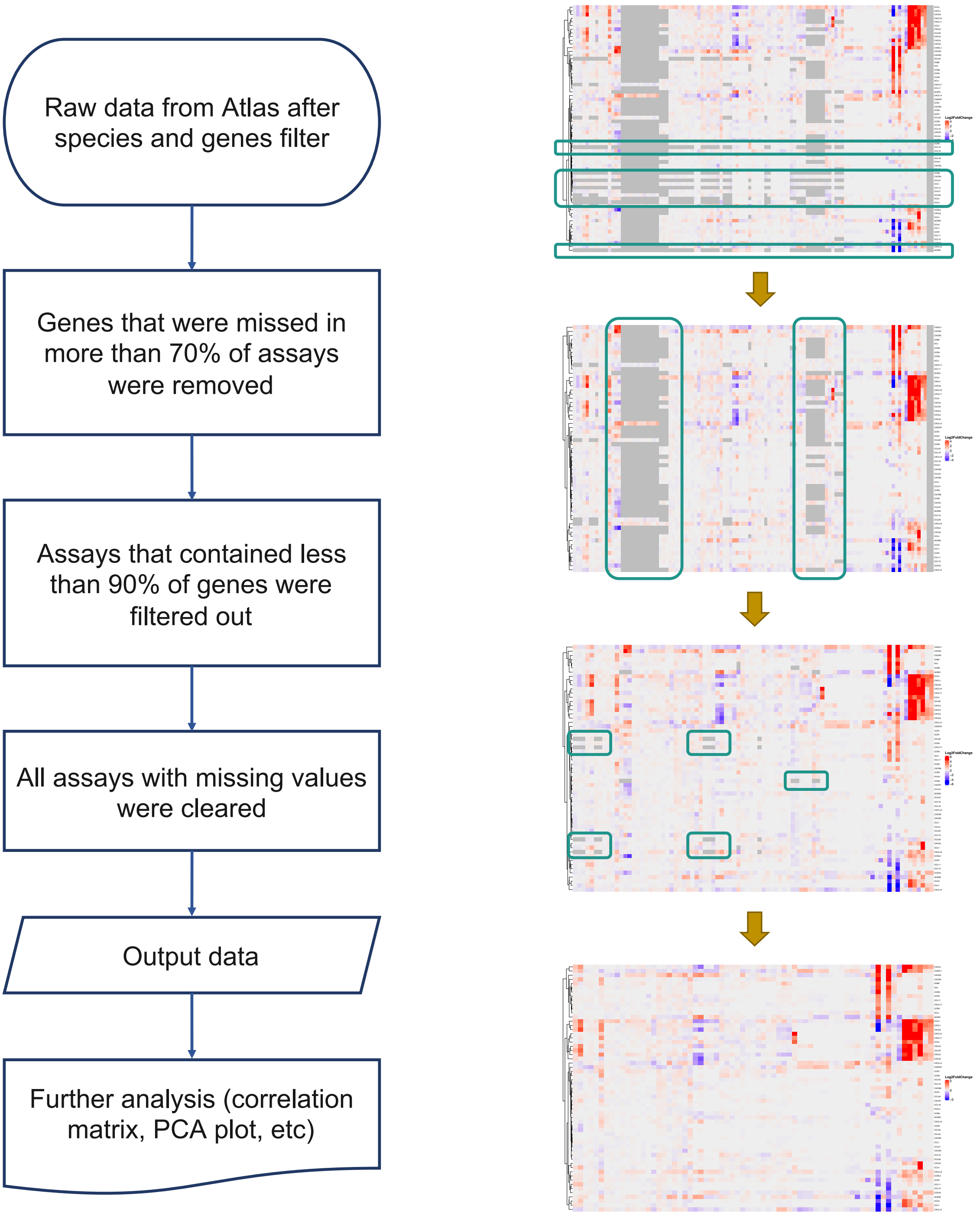


**Supplementary Figure 1. Schematic explanation of data analysis from the EMBL-ELI expression atlas.**

Supplementary Figure 2

### Human

#### Brain

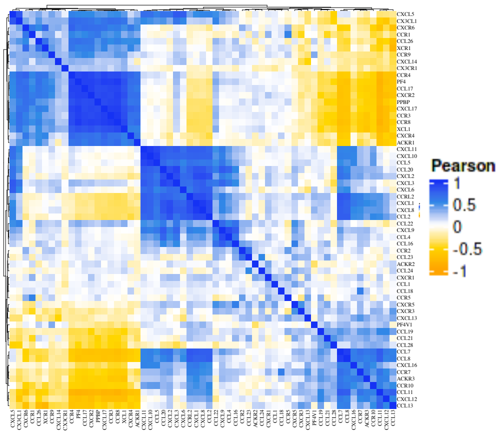


#### Lungs

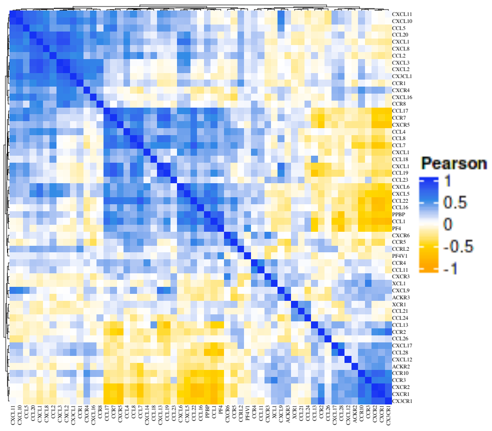


#### Lymph node

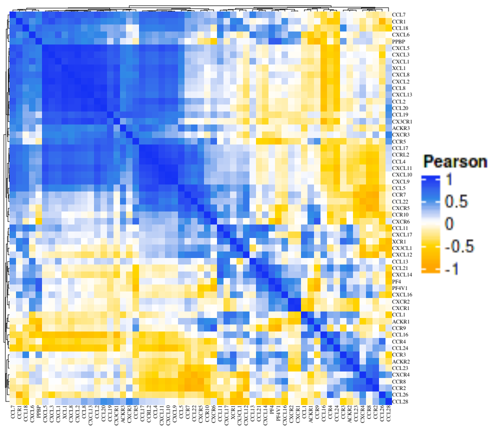

**Supplementary Figure 2. Human tissues have tissue specific chemokine ligand and receptor expression patterns.** EMBL-ELI expression atlas heat map analysis of chemokine ligands and receptors across human brain, lungs and lymph node.

Supplementary Figure 3

### Mouse

#### All tissues

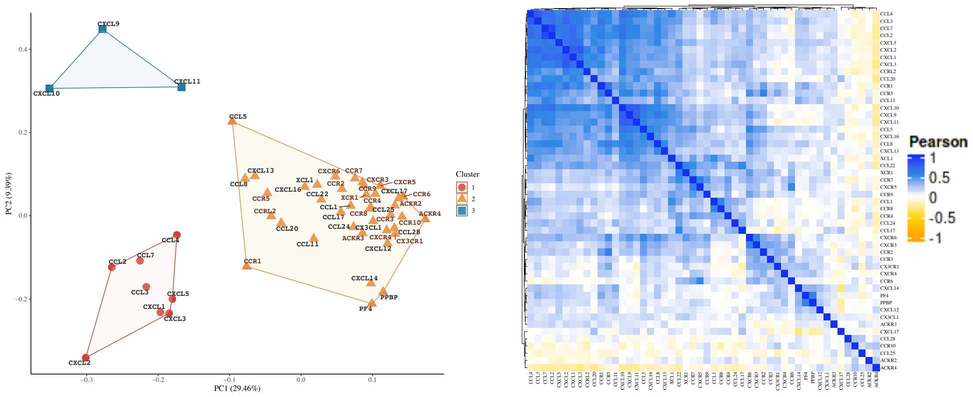


#### Brain

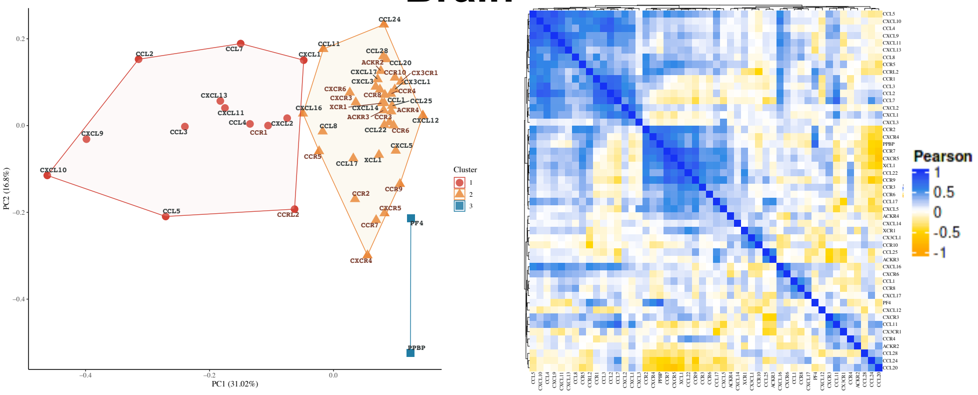


#### Lungs

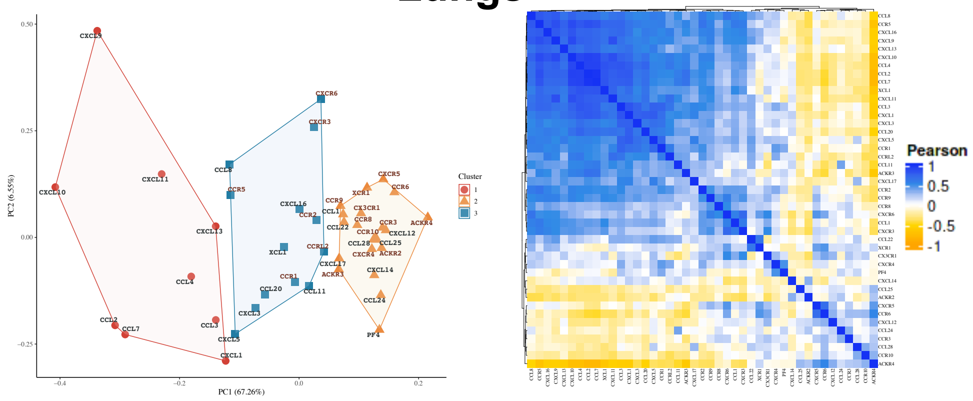


#### Lymph node

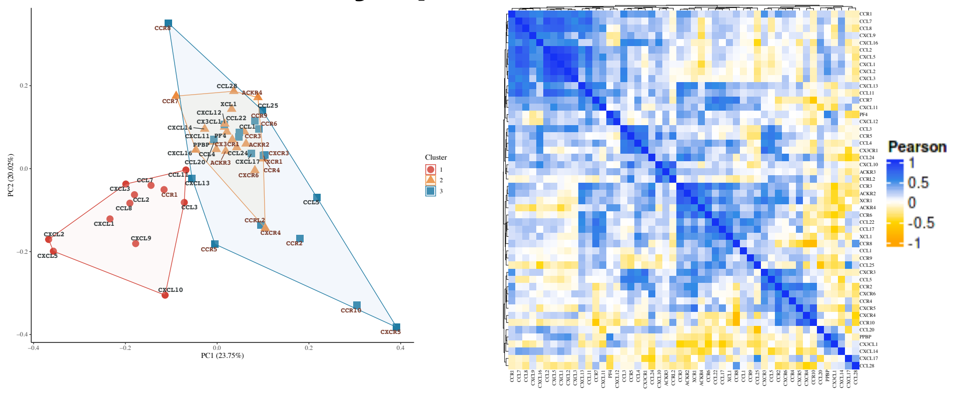

**Supplementary Figure 3. Mouse tissues have tissue specific chemokine ligand and receptor expression patterns.** EMBL-ELI expression atlas PCA and heat map analysis of chemokine ligands and receptors across mouse pooled tissues, mouse brain, lungs and lymph node.

Supplementary Figure 4

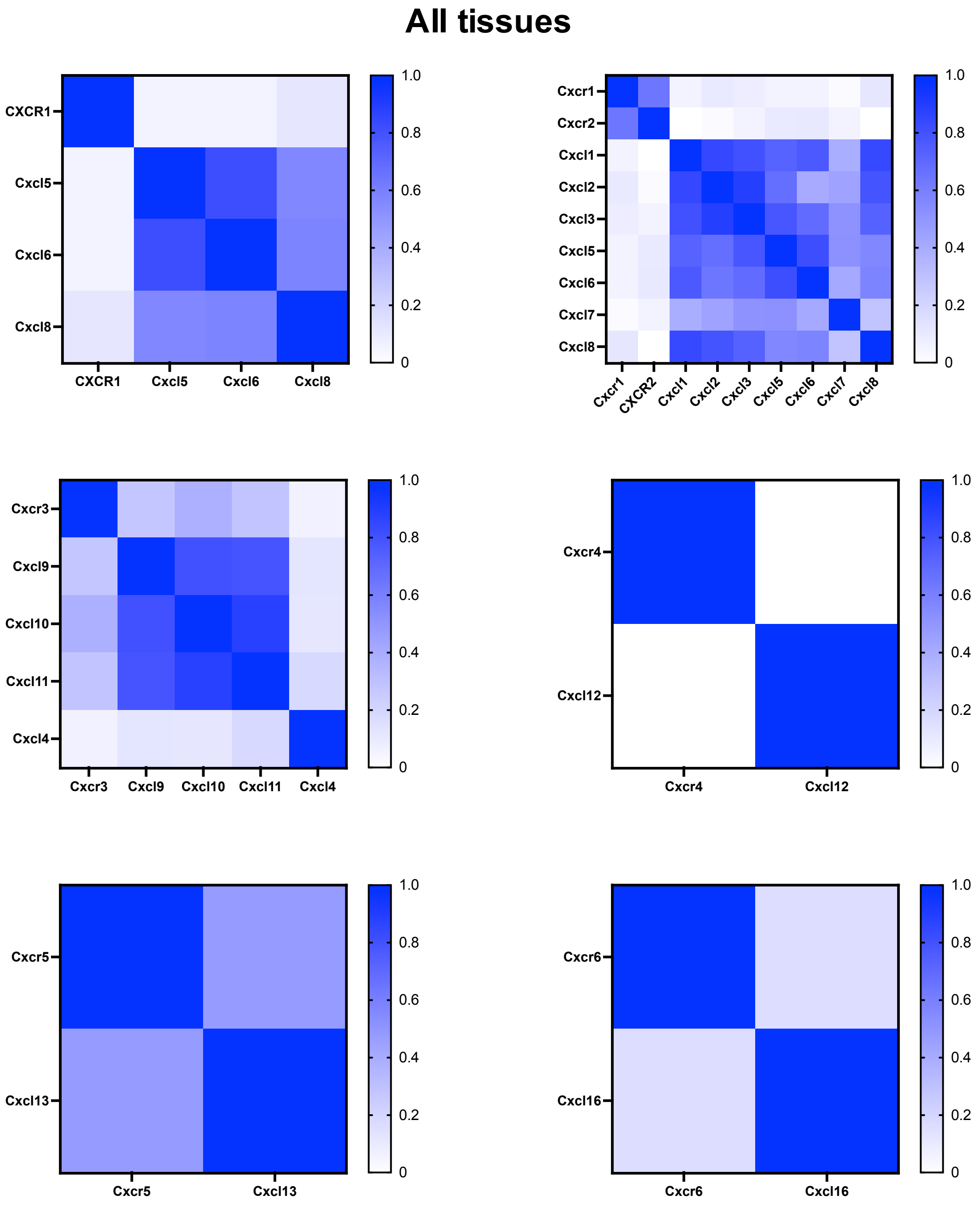

**Supplementary Figure 4. Chemokine receptors have different transcriptional relationships with their ligands.** EMBL-ELI expression atlas heat map analysis of Pearsons correlation between CXCR1, 2, 3, 4, 5 and 5 and their individual ligands.

Supplementary Figure 5

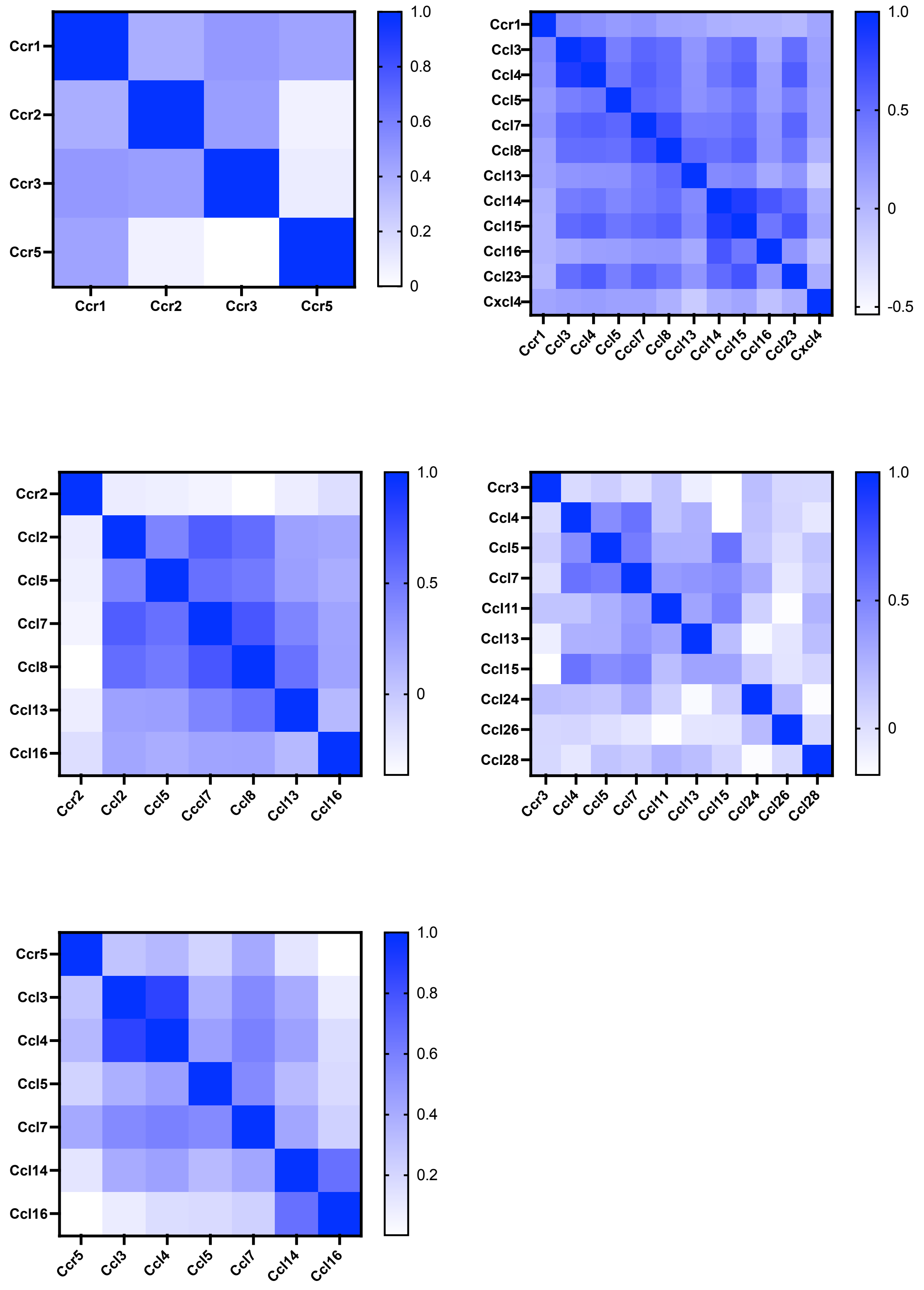

**Supplementary Figure 5. Chemokine receptors have different transcriptional relationships with their ligands.** EMBL-ELI expression atlas heat map analysis of Pearsons correlation between CCR1, 2, 3 and 5 and their individual ligands.

#### Supplementary Figure 6

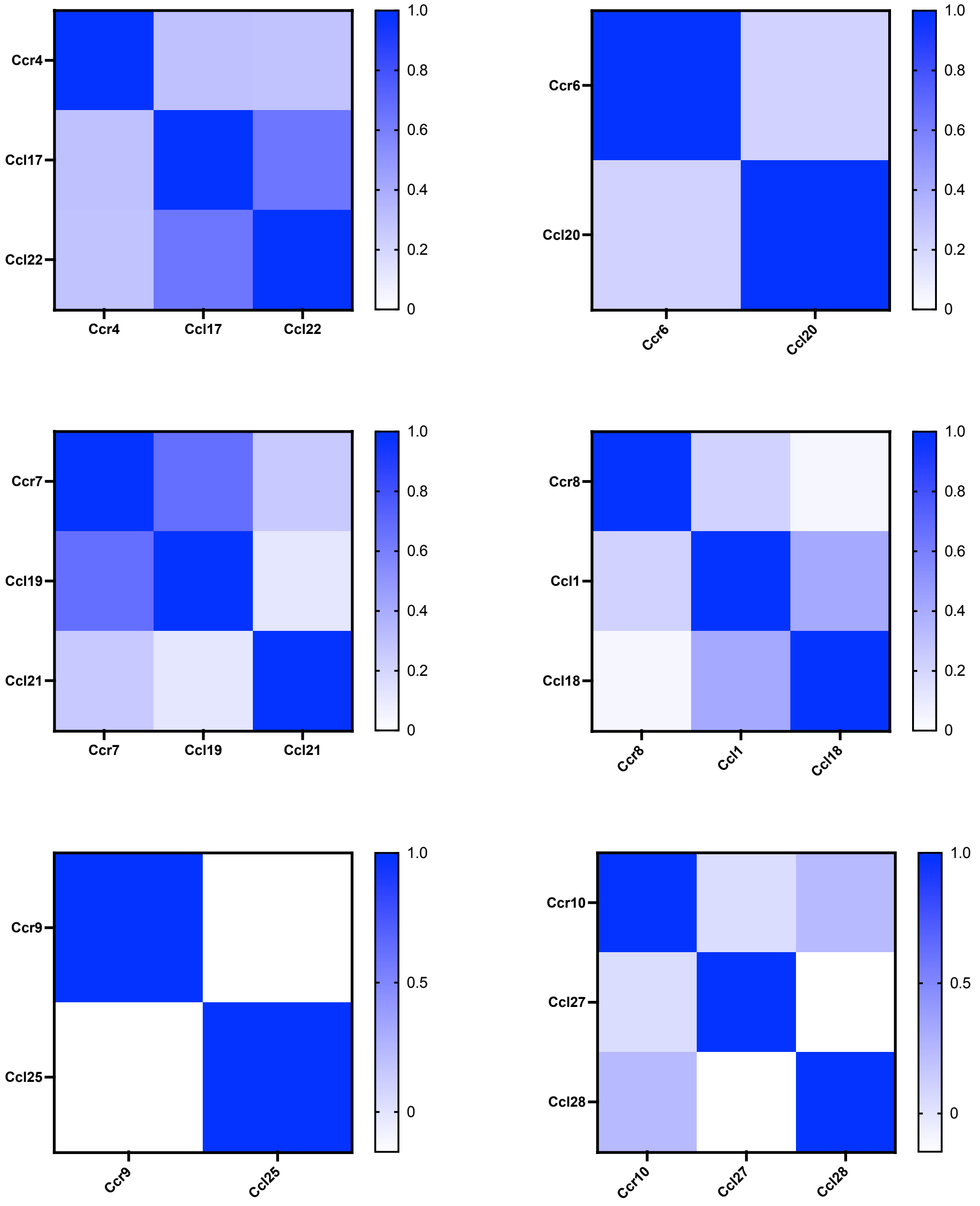

**Supplementary Figure 6. Chemokine receptors have different transcriptional relationships with their ligands.** EMBL-ELI expression atlas heat map analysis of Pearsons correlation between CCR4, 6, 7, 8, 9 and 10 and their individual ligands.

Supplementary Figure 7

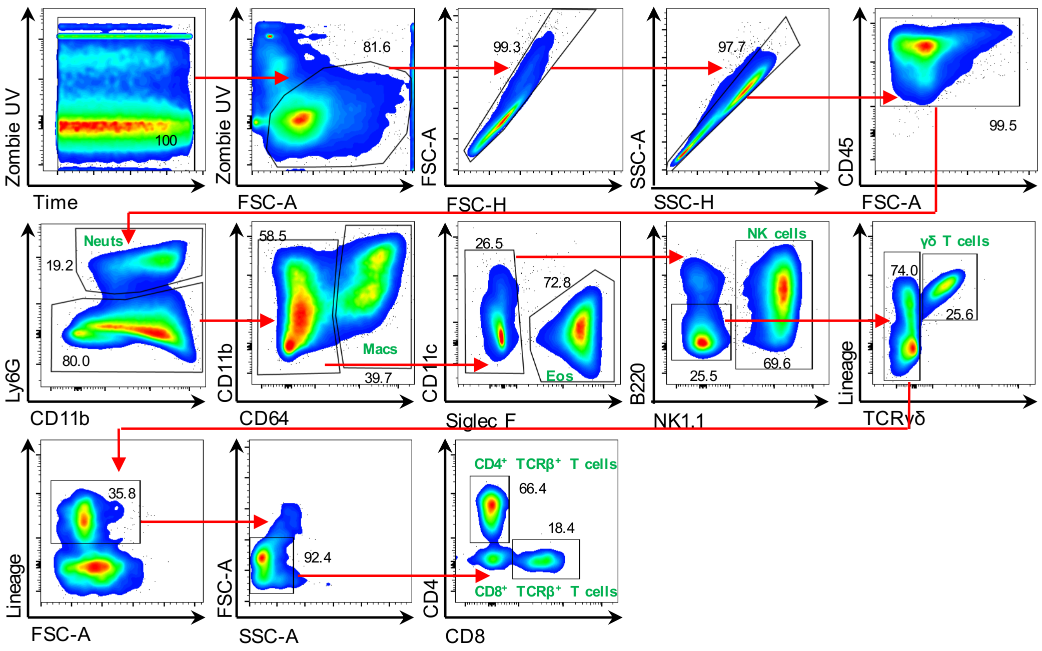


**Supplementary Figure 7. Flow cytometric gating strategy for identyfying air pouch fluid derived immune cells.** Cells isolated from air pouch fluid were stained and cell populations (highlighted in green) were defined based on their expression profiles of the surface markers stated. Plots are representative of >3 experiments although the proportions of different cell types differed. Numbers adjacent to boxed areas indicate the percentages of cells that fall within each gate.

### Supplementary Figure 8

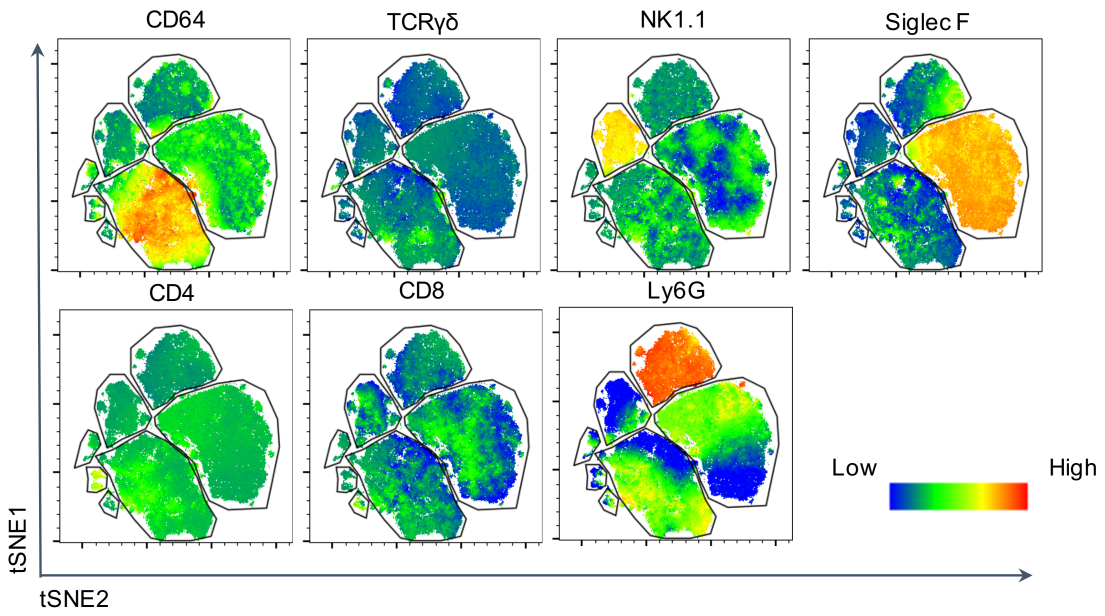


**Supplementary Figure 8. tSNE analysis of cell populations flushed from the air pouch after chemokine injection.** Representative tSNE analysis of cells flushed from the chemokine stimulated air pouch for expression oft he indicated flow cytometry markers.

Supplementary Figure 9

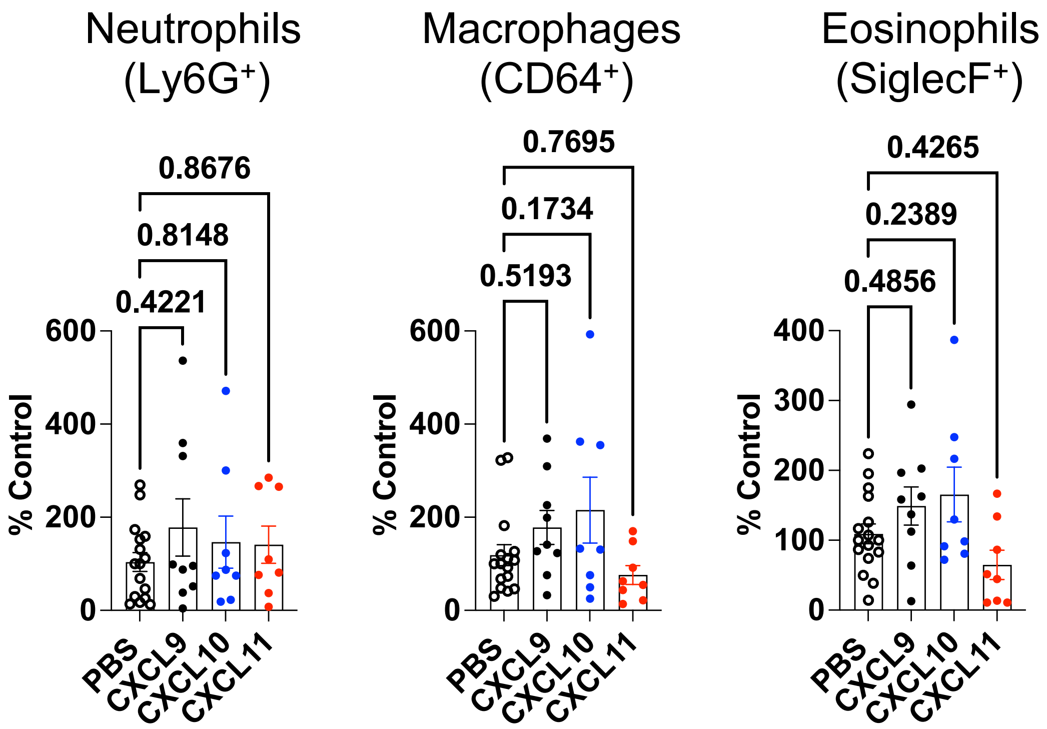


**Supplementary Figure 9. CXCR3 ligands do not recruit neutrophils, macrophages or eosinophils to the air pouch.** CXCL9, 10 and 11 were injected into the air pouch and 24 hrs later cells were analysed later to quantify the number of recruited neutrophils, macrophages or eosinophils.

### Supplementary Figure 10

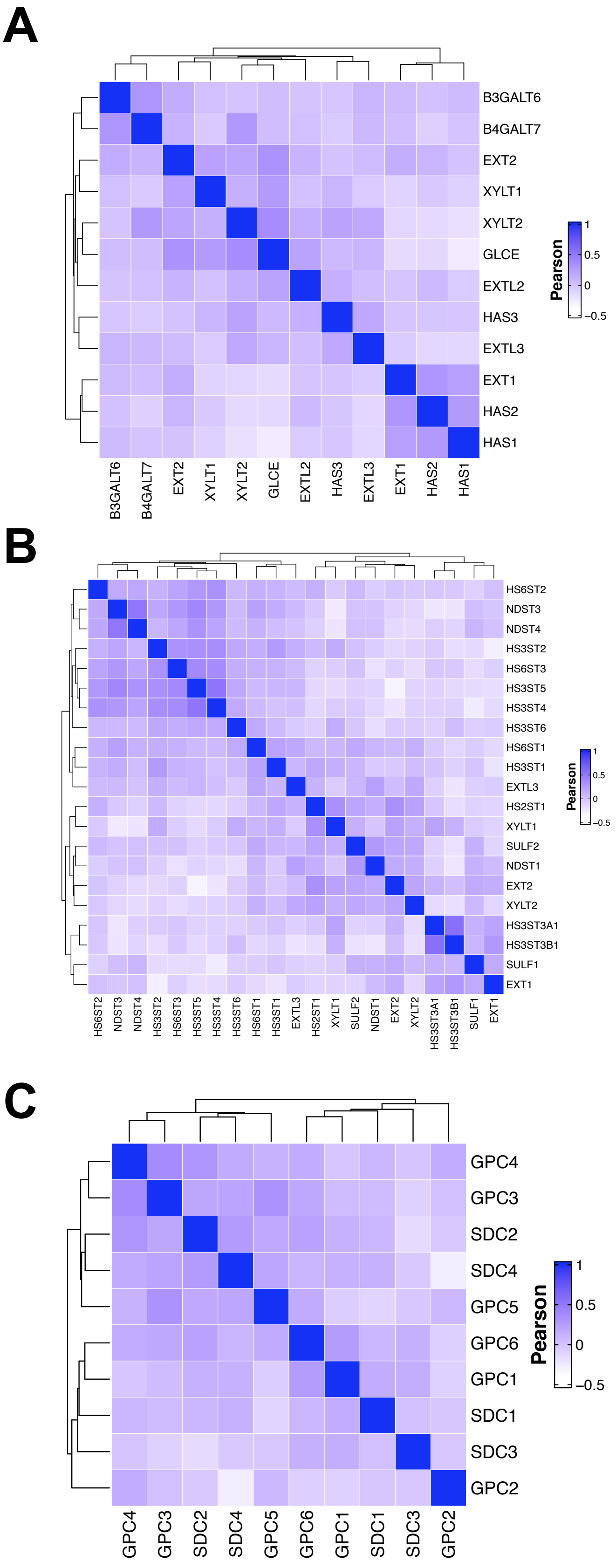

**Supplementary Figure 10. ECM proteoglycan GAG synthesis, sulphation and protein core genes show little transcriptional correlation.** EMBL-ELI expression atlas heat map analysis of Pearsons correlation between genes that mediate (A) GAG chains synthesis, (B) GAG chain sulphation or (C) proteoglycan protein core production.

Supplementary Figure 11

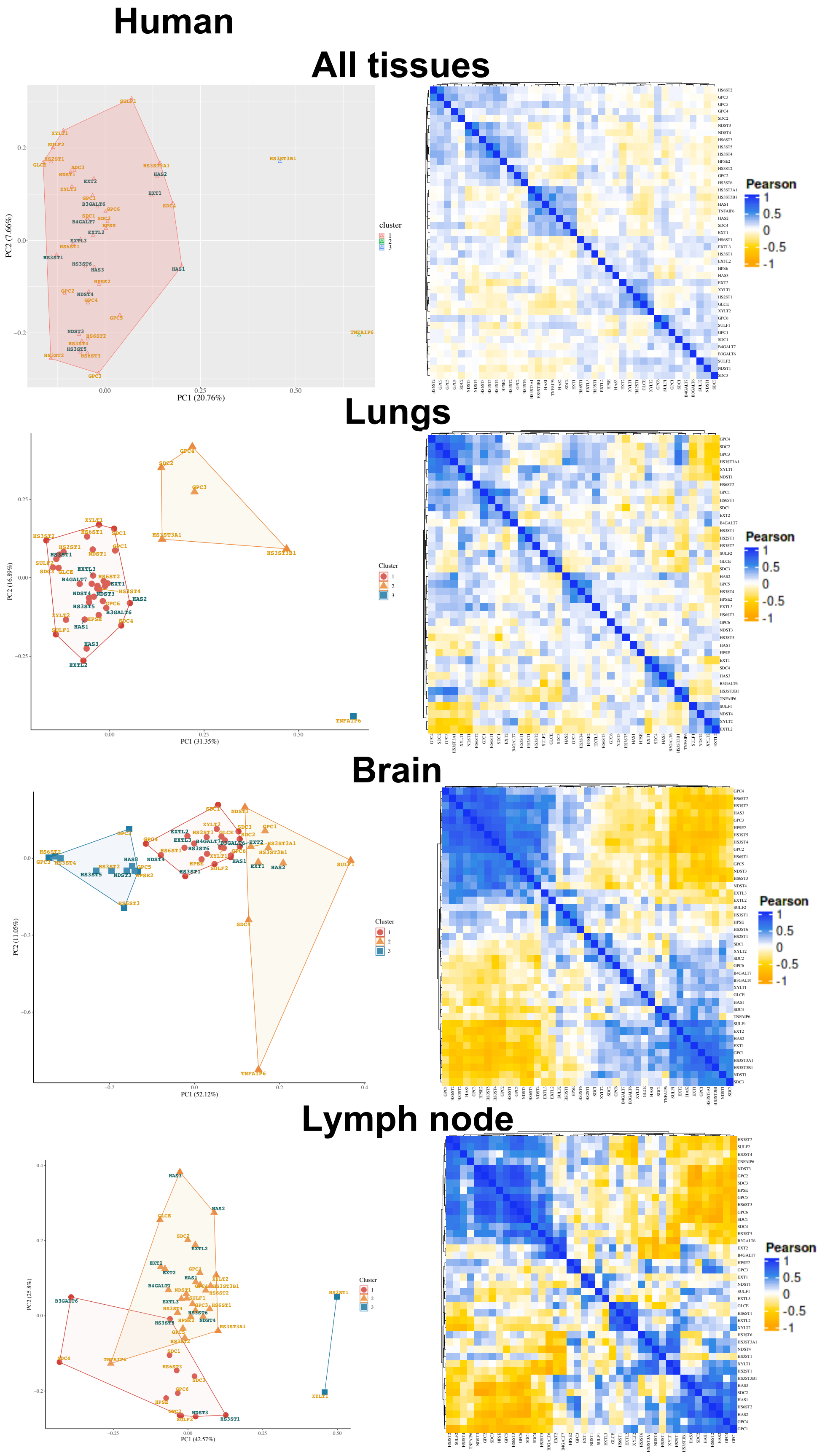

**Supplementary Figure 11. Human ECM GAG genes have tissue specific correlation signatures.** EMBL-ELI expression atlas PCA and heat map analysis of Pearsons correlation between genes involved in ECM GAG synthesis and modification across human tissues.

Supplementary Figure 12

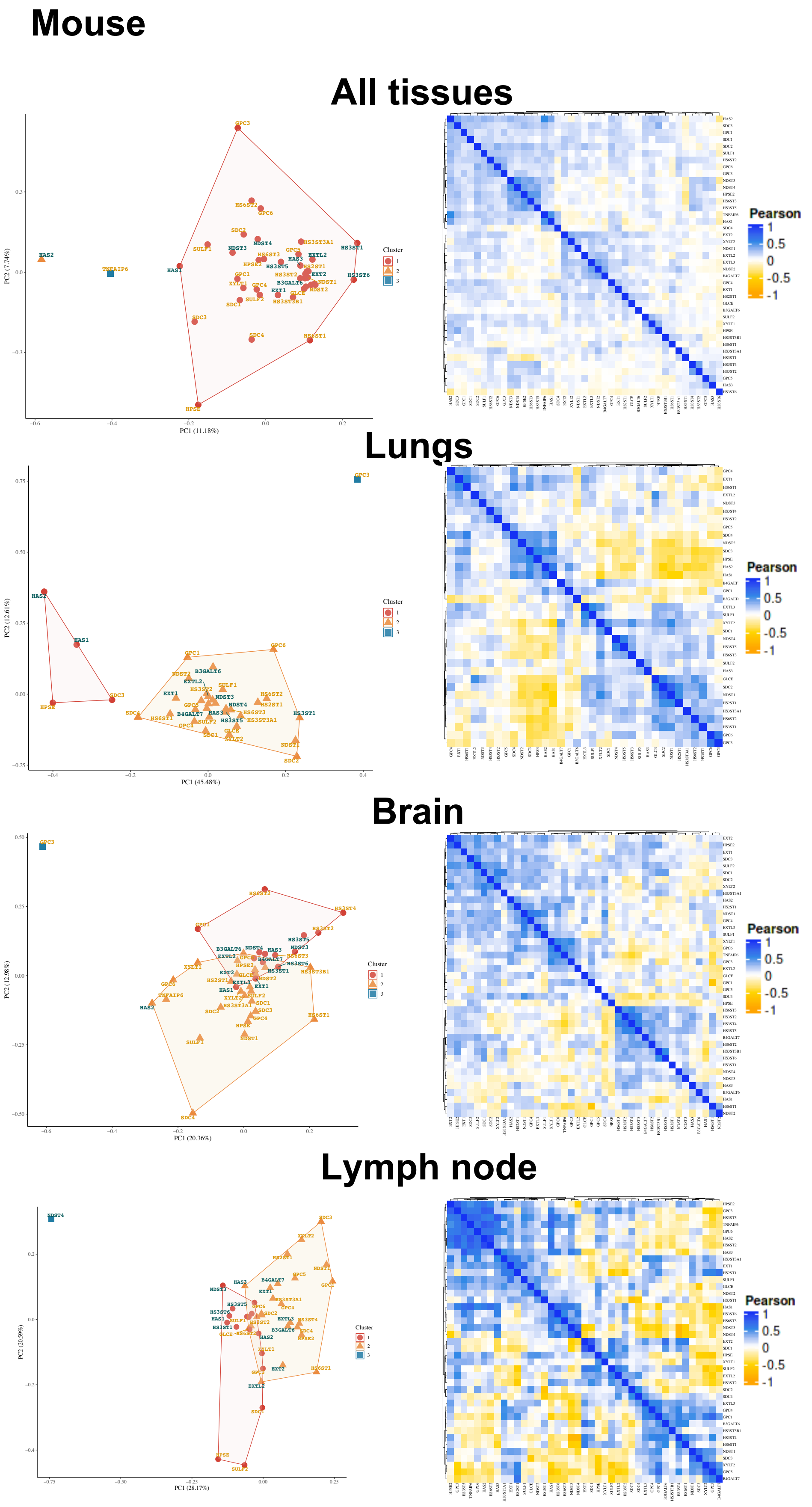

**Supplementary Figure 12. Mouse ECM GAG genes have tissue specific correlation signatures.** EMBL-ELI expression atlas PCA and heat map analysis of Pearsons correlation between genes involved in ECM GAG synthesis and modification across murine tissues.
